# Systematic analysis of the genomic features involved in the binding preferences of transcription factors

**DOI:** 10.1101/2022.08.16.504098

**Authors:** Raphaël Romero, Christophe Menichelli, Jean-Michel Marin, Sophie Lèbre, Charles-Henri Lecellier, Laurent Bréhélin

**Affiliations:** LIRMM, Univ Montpellier, CNRS, Montpellier, France; IMAG, Univ. Montpellier, CNRS, Montpellier, France; Institut de Génétique Moléculaire de Montpellier, University of Montpellier, CNRS, Montpellier, France; Univ. Paul-Valéry-Montpellier, Montpellier, France

## Abstract

Transcription factors (TFs) orchestrate gene expression and are at the core of cell-specific phenotypes and functions. One given TF can therefore have different binding sites depending on cell type and conditions. However, the TF core motif, as represented by Position Weight Matrix for instance, are often, if not invariably, cell agnostic. Likewise, paralogous TFs recognize very similar motifs while binding different genomic regions. We propose a machine learning approach called TFscope aimed at identifying the DNA features explaining the binding differences observed between two ChIP-seq experiments targeting either the same TF in two cell types or treatments or two paralogous TFs. TFscope systematically investigates differences in i) core motif, ii) nucleotide environment around the binding site and iii) presence and location of co-factor motifs. It provides the main DNA features that have been detected, and the contribution of each of these features to explain the binding differences. TFscope has been applied to more than 350 pairs of ChIP-seq. Our experiments showed that the approach is accurate and that the genomic features distinguishing TF binding in two different settings vary according to the TFs considered and/or the conditions. Several samples are presented and discussed to illustrate these findings. For TFs in different cell types or with different treatments, co-factors and nucleotide environment often explain most of the binding-site differences, while for paralogous TFs, subtle differences in the core motif seem to be the main reason for the observed differences in our experiments.

The source code (python), data and results of the experiments described in this article are available at https://gite.lirmm.fr/rromero/tfscope.

## Introduction

The programming of gene expression is the primary mechanism that controls cellular phenotype and function. At the DNA level, transcription factors (TFs) are supposed to play a key role in this control. These proteins bind DNA sequence through specialized DNA binding domains (DBDs) to enhance or repress the transcription of their target genes. DBDs bind preferentially to specific DNA sequences, which are resumed in statistical models known as Position Weight Matrices (PWMs) [49]. Most of the times, PWMs are obtained from dedicated probabilistic models—the Position Probability Matrices (PPMs)—, which are available for many TFs in databases like JASPAR [15] and HOCOMOCO [28]. PWMs can be used to compute binding affinities and to identify potential binding sites in genomes. However, contrary to bacterial DBDs which recognize sequences that often have sufficient information content to target particular genomic positions, most eukaryotic DBDs recognize short binding motifs (around 10bp) that are not sufficient for specific targeting in the usually large (*e*.*g*. 10^9^bp) eukaryotic genomes [52]. This purely statistical analysis has been corroborated by genome-wide studies based on sequencing approaches (ChIP-seq, ChIP-exo, CUT&RUN) that have been applied to hundreds of TFs in order to determine their binding profiles in various cell types and conditions [13]. These studies showed that most TFs only associate with a small subset of their potential genomic sites *in vivo* [48], and that the binding sites of a given TF often vary substantially between cell types and conditions [44]. Furthermore, as the number of DBD families in a genome is small with regard to the number of TFs, TFs paralogs from the same DBD family often share very similar binding motifs, yet they usually show distinct binding sites *in vivo* [21, 42, 27]. Thus, it is now evident that DBD motifs as resumed by PWMs are not sufficient to completely determine TF binding in a specific cell or condition. On the other hand, several studies revealed that a substantial part of the *in vivo* binding sites lack an obvious match with the known binding motif of the target TF [48, 27].

At this point, it is important to emphasize the strong links that exist between TF binding and histone marks [14]. Also, ChIP-seq experiments revealed that most TF binding sites (TFBSs) lie within highly accessible (*i*.*e*., nucleosome-depleted) DNA regions [45]. However, it remains unclear whether these chromatin states are a cause or a consequence of TF binding [20]. Moreover, recent approaches based on machine learning, and specifically convolutional neural networks (CNNs), have shown that transcription factor binding but also gene expression as well as histone modifications and DNase I-hypersensitive sites can be predicted from DNA sequence only, often with surprisingly high accuracy [50, 54, 36, 24, 47, 2]. The good predictive performances of these approaches suggest that a large part of the instructions for gene regulation and TF binding lies at the level of the DNA sequence.

Several mechanisms based on specific DNA features have thus been proposed to complement DBD motifs and to explain how TFs target precise genome locations. The current view is that TF combinations underlie the specificity of eukaryotic gene expression regulation [11], with several TFs competing and collaborating to regulate common target genes. Multiple mechanisms can lead to TF cooperation [34, 37]. In its simplest form, cooperation involves direct TF-TF interactions before any DNA binding. But cooperation can also be mediated through DNA, either with DNA providing additional stability to a TF-TF interaction [22], or without any direct protein-protein interaction, as in the pioneer/settler hierarchy described in Sherwood et al. [43] or in a non-hierarchical cooperative system such as the billboard model for enhancers [3, 33].

Besides TF combination, other studies have investigated the role the genomic environment around TFBS may have on binding specificity, revealing that some TFs have a preferential nucleotide content in the flanking positions of their core binding sites [30, 12]. Other studies have proposed that much larger regions containing repetitive sequences or multiple occurrences of low-affinity motifs may play an active role in TF binding [1, 9, 27]. Finally, another possibility that may be underestimated and that could also explain binding specificity in certain cases is that, depending on cell, condition, or TF paralog, the binding motif may actually differ, showing globally the same PWM to our eyes, but slightly differing on specific positions.

All these mechanisms have been independently studied on specific cases, but a global computational approach is still missing to investigate their role and relative importance in an automatized manner. The above-mentioned deep learning approaches are able to capture and combine the different DNA features involved, but identifying them from the CNNs remains a difficult task [26, 17]. Although interesting methods are being developed to post-analyze CNN predictions and to identify single nucleotides and motifs (see *e*.*g*. [54, 4, 26]), disentangling all mechanisms/features captured by a CNN remains unreliable.

Here, we propose a machine learning approach called TFscope specially designed to explain the binding differences observed between two settings: two cell types, two treatments, or two paralogous TFs. Our method directly compares the two ChIP-seq data associated with the two settings, by considering only regions bound either in one or the other experiment. This strategy has two advantages. First, by focusing on the binding differences, we obviously gain sensitivity for identifying the genomic features that best explain these differences. Second, we circumvent the common problem of the background definition which arises in all studies that aim to distinguish bound (foreground) versus unbound (background) genomic regions in a given cell type. While the definition of the foreground is straightforward, the definition of the background is often much more challenging and strongly influences the results and the conclusions (see for example references [51, 50, 35, 53] for interesting considerations about the background issue).

Given two ChIP-seq data, our method systematically investigates the importance of i) the core motif, ii) the genomic environment and iii) the cooperative TFs for predicting the binding differences between two data. TFscope is based on three different modules that capture these three levels of information. The first module captures the potential differences in the core motif. This module is based on a new method that learns discriminative PWMs. It is worth noting that well known approaches such as DREME/STREME [5], DAMO [39], Homer [19], etc. have been already proposed for this task. These methods are however designed for a slightly different and computationally more complex problem, that is not exactly the same as ours. As a consequence, they rely on sub-optimal heuristics while an optimal algorithm exists for our problem. The second module captures the nucleotidic environment in the form of short k-mers (2-4 bps) enriched in specific regions around the core motif and is based on our DExTER method [32]. The third module is a refinement of our TFcoop method that identifies co-factors and TF combinations involved in the binding of a target TF [47]. In a final step, these data are used together in a global predictive model that is used to quantify the relative importance of each information for the problem at hand. Hence, in contrast to CNN based methods [53, 4], our approach completely controls the predictive features inputted into the model. This allows to easily measure the importance of each feature by computing the loss of accuracy induced by its withdrawal from the model, something very challenging to do with classical CNN approaches.

We applied TFscope to more than 350 pairs of ChIP-seq targeting either a common TF in two different cell types or treatments, or two paralogous TFs in the same cell type. Our results showed that classification is very often accurate and that the most important DNA features greatly vary depending on TFs and conditions. For TFs in different cell types or with different treatments, either co-factors or the nucleotidic environment often explainsmost of the binding-site differences. Moreover, when co-factors are involved, which is the most frequent case, their position on the DNA relative to the core motif is also important. On the contrary, for paralogous TFs the core motif seems to be the most important factor in our experiments. Although the motifs of paralogous TFs show very similar PWMs, subtle differences at specific positions explain most of the binding differences.

## Results

### TFscope overview

TFscope aims to identify the genomic features responsible for the binding differences observed between two ChIP-seq experiments. Typically, TFscope can be used to identify the differences between two experiments targeting the same TF in different cell types or conditions, or two experiments targeting two paralogous TFs that share similar motifs. TFscope takes in input two sets of ChIP-seq peaks corresponding to the two ChIP-seq experiments and then runs the three steps illustrated on Figure 1: sequence selection & alignment, feature extraction, and model learning. In the sequence alignment step, TFscope first identifies the peaks that are unique either to the first or the second set (see Material and Methods). All common peaks are discarded for the analysis (this point is discussed in the experiments below). Then, TFscope identifies the most likely binding site using a strategy similar to Centrimo [6] and UniBind [16], and parses the sequence around the peak summit with the PWM associated with the target TF (if several versions of the motif are available or if the analysis involves two paralogous TFs with similar motifs, the most discriminative PWM is chosen to scan the two sets of sequences; see Methods). The FIMO tool [18] is used for this analysis, and the position with the highest PWM score is used as an anchor point to extract the 1Kb long sequence centered around this position. At the end of the alignment step, we get two classes of sequences centered on the most likely TFBSs of the ChIP-seq peaks given in input. Sequences with no occurrence of the motif around the peak summit are discarded.

**Figure 1.**
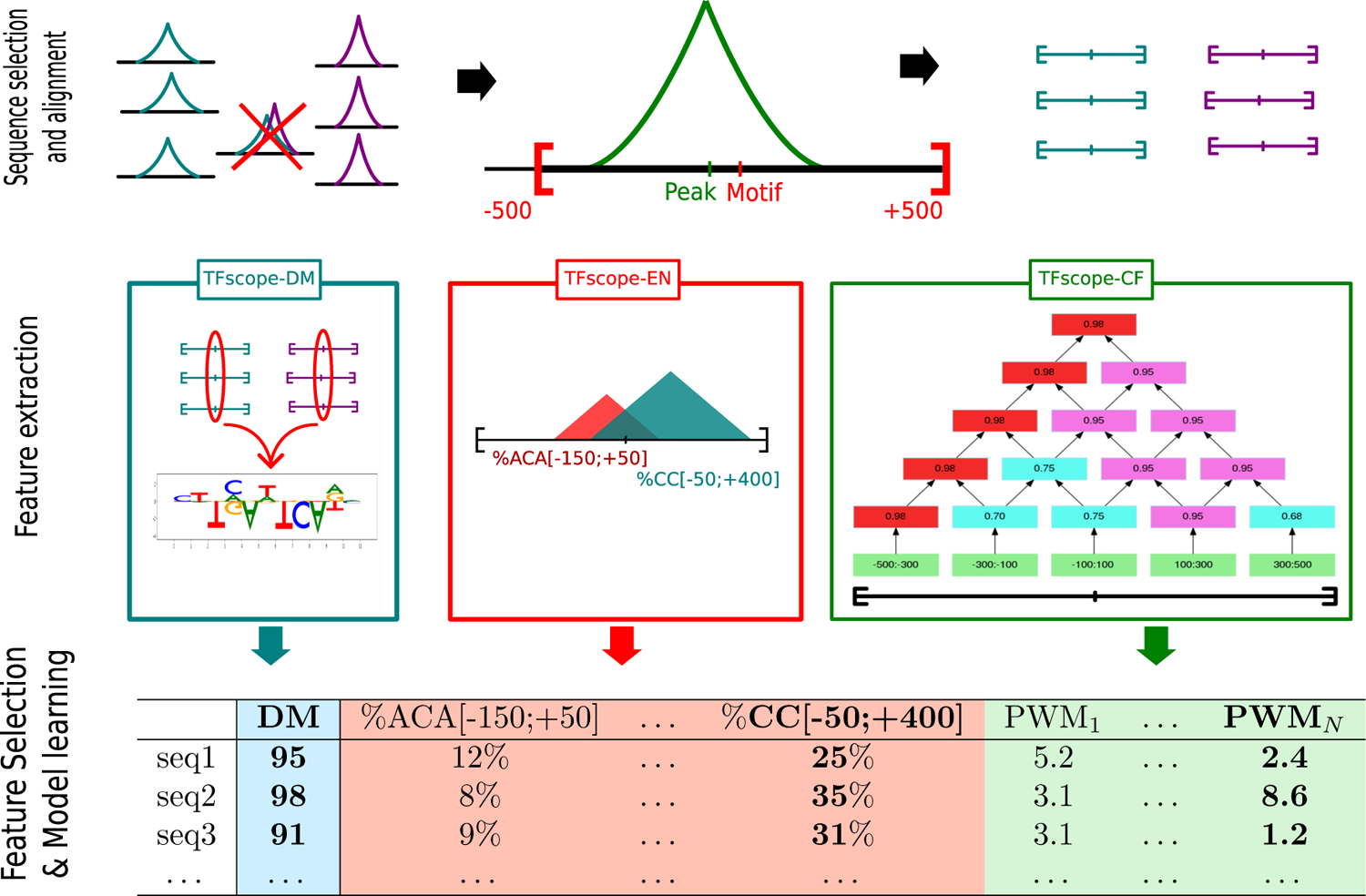
The TFscope approach. In the first step (sequence selection and alignment), peaks associated with both ChIP-seq experiments are removed. The most likely TFBSs of the remaining peaks are identified and used to extract the 1Kb sequences centered on these sites. All sequences are then used for the second step (feature extraction). Three dedicated modules extract three kinds of genomic features that can be useful for discriminating the two classes. The TFscope-DM module learns a new PWM that can discriminate the sequences on the basis of the core motif solely. The TFscope-EN module searches for specific nucleotidic environments (*i*.*e*. frequency of specific k-mers in specific regions) that are different in the two classes. The TFscope-CF module searches for binding sites of specific co-factors whose presence in specific regions differs between the two classes. All these features (variables) are then gathered into a long table, and a logistic model (Expression (1)) is learned on the basis of these data (Feature selection and model learning). A special penalty function (LASSO) is used during training, for selecting only the best variables in the model (in bold in the table).

TFscope then runs three modules detailed below to extract three kinds of DNA features that are discriminative of the two sequence classes (feature-extraction step). The first module (TFscope-DM) learns a new PWM of the core motif (see below). This PWM is different from the original PWM used to parse the sequence, as it focuses on the potential differences of core motif that may exist between the two sequence sets. This module returns a single variable DM(*s*), which is the score of the new PWM on each sequence *s*. The second module (TFscope-NE) searches for pairs of (k-mer,region) for which the frequency of the k-mer in the defined region is different between the two sets of sequences. For example, we may observe that the frequency of the 3-mer ACA in region [−150 : +500] (0 being the anchor point of the sequences) is globally higher in sequences of the first class than in those of the second class. The idea is to capture the differences in nucleotidic environment that may exist between the two classes. We use for this module a slight modification of the DExTER method recently proposed to identify long regulatory elements [32]. This module returns a potentially large set of variables NE_*i*_(*s*) that corresponds to the frequency of *i*th k-mer in the associated *i*th region for each sequence *s*. The third module (TFscope-CF) uses a library of PWMs (in the experiments below the JASPAR2020 library is used [15]) and searches for pairs of (PWM,region) for which the score of the PWM in the identified region is different between the two sets of sequence (see below). The idea is to identify all co-factors of the target TF whose binding sites differ between the two classes: either because these binding sites are in majority present in one class and not the other, or because the locations of these binding sites are different between the two classes. This third module returns a set of variables CF_*j*_(*s*) that corresponds to the score of the *j*th PWM in the identified *j*th region for each sequence *s*.

All variables are then integrated into a global model that aims to predict if a sequence belongs to the first or the second class (learning step). We used a logistic regression model:

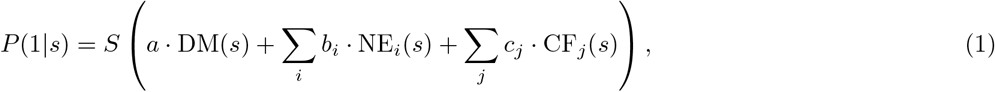

where *P* (1|*s*) is the probability that sequence *s* belongs to the first class, *S* is the sigmoid function, DM(*s*) is the score of the discriminative motif for sequence *s*, NE_*i*_(*s*) is the value of the *i*th nucleotidic-environment variable for sequence *s*, CF_*j*_(*s*) is the value of the *j*th co-factor variable for sequence *s*, and *a, b*_*i*_ and *c*_*j*_ are the regression coefficients which constitute the parameters of the model. Because the set of variables identified by the last two modules is usually large and variables are often correlated, the model is trained with a LASSO penalty function [46] that selects the most relevant variables—*i*.*e*. many regression coefficients (*a, b*_*i*_ and *c*_*j*_) are set to zero. Finally, once a model has been trained, its accuracy is evaluated by computing the area under the ROC (AUROC) on several hundred sequences. To avoid any bias, this is done on a set of sequences that have not been used in the previous steps.

### TFscope-DM: Identification of differences in the core motif

The first TFscope module learns a new discriminative PWM. Recall that at the end of the alignment step, the most likely binding site of each ChIP-seq peak has been identified with the JASPAR PWM associated with the TF, and all sequences are aligned on these sites. If several versions of the PWM are available, the most discriminative PWM is used (see Methods). We then extract the *K*-length sub-sequence corresponding to the occurrence of the motif in each sequence (*K* being the size of the PWM). The first module aims to learn a new PWM that could discriminate these two sets of *K*-length sequences. First, each sequence *s* is one-hot encoded in a *K* × 4 matrix **s**. Then, a logistic model with *K* × 4 parameters is learned to discriminate the two classes of sequences:

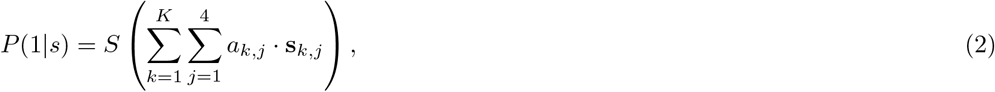

with *P* (1 | *s*) the probability that sub-sequence *s* belong to the first class, *S* the sigmoid function, **s**_*k,j*_ the entry of the one-hot matrix **s** indicating whether the *k*th nucleotide of sequence *s* corresponds to the *j*th nucleotide of {*A, T, G, C*} or not, and *a*_*k,j*_ the regression coefficients of the model.

Once this model has been learned, it can be used to predict if a sequence belongs to the first or the second class. The sigmoid function being monotonically increasing, this can be done easily by computing the linear function inside the parenthesis of Expression (2) and using the result as a score reflecting the likelihood of class 1. Interestingly, this score function has exactly the same form as the one used to compute a score with a PWM (see Material and methods). As a consequence, the logistic model of Expression (2) is strictly speaking a regular PWM with parameters *a*_*k,j*_. The interest to learn a PWM in this way is two folds. First, we take advantage of all the algorithmic and theory developed for logistic regression. Most notably, as the likelihood function of a logistic model is convex, we have the guarantee that the learned model is optimal, which means that the inferred discriminative PWM is the best PWM for our problem. This is an important difference from the approaches already proposed to learn a discriminative PWM, such as DAMO [39] or STREME [5]. The reason for this is that these approaches do not exactly address the same problem as ours: they do not search for a PWM that discriminates two sets of sequences perfectly aligned and of the same length as the PWM. Instead, they take as input two sets of sequences usually much longer than the PWM, and their goal is to identify a motif whose presence can be used to discriminate the two sets, a problem known to be NP-hard [31]. As a consequence, these approaches rely on heuristics and do not warrant returning the best PWM for our problem (see section Discussion for more details on these differences). The second advantage to learn a PWM via a logistic regression approach is that we can include a LASSO penalty in the optimization procedure in order to obtain a model with fewer variables [46] (see Material and methods). In practice, this means that many parameters *a*_*k,j*_ are set to zero, and hence that the resulting PWM is simpler and easier to interpret.

It is important to note that, as DAMO [39], the PWMs output by our method are not obtained from position probability matrices (PPMs), which are the probabilistic models that are often associated with PWMs. This avoids the constraints attached to PPMs (see section Discussion and the work of Ruan and Stormo [38] for more details) but this also impedes to represent PWMs with the classical logo graphics based on information theory [40]. Instead, our PWMs are represented by “mirror-logos” such as the one on Figure 2B (middle). These logos provide the sign of the parameters, which allows to easily distinguish the nucleotides that are more present in sequences of one or the other class.

**Figure 2.**
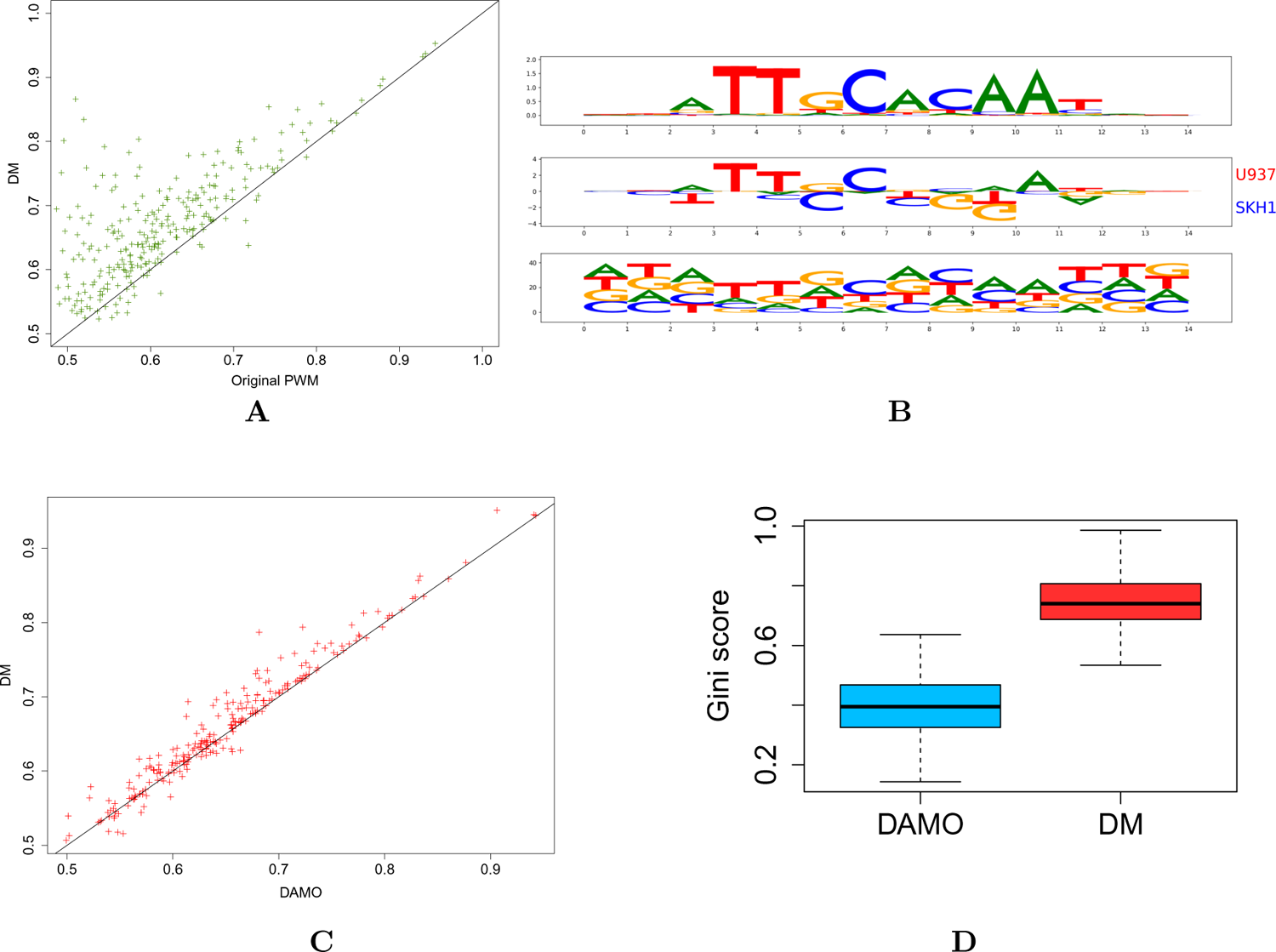
TFscope learns discriminative and informative motifs. **A** AUROCs achieved by the TFscope PWMs *vs*. original PWMs on the 272 experiments. **B(up)** Original (JASPAR) PWM of TF USF2. **B(middle)** discriminative PWM learned by TFscope for discriminating USF2 binding between HepG2 and GM12878. **B(bottom)** discriminative PWM learned by DAMO on the same training set. **C** AUROCs achieved by the TFscope PWMs *vs*. DAMO PWMs on the 272 experiments.**D** Gini score of the PWMs learned by TFscope and DAMO. The higher the Gini score, the simpler the model.

### TFscope-NE: Identification of differences in nucleotidic environment

The second TFscope module extracts features related to the nucleotidic environment around the core binding motif. More precisely, this module constructs variables defined by a pair (kmer,region) such that the frequency of the identified k-mer in the identified region is, on average, different in the two classes. We used for this a slight modification of the DExTER method initially proposed to identify pairs of (kmer,region) whose values are correlated with an expression signal. The optimization function of DExTER has thus been modified to return variables correlated with classes rather than with expression signal (see Material and method). The TFscope-NE module explores short k-mers up to length 4. To prevent this module to capture information related to the core-motif, this motif is masked before running the TFscope-NE analysis.

### TFscope-CF: Identification of differences in co-factor combinations

The third TFscope module extracts features related to co-factors. This module constructs variables defined by a pair (PWM,region) such that the score of the PWM in the identified region is, on average, different between the two classes. For example, one can observe that sequences of the first class often have a potential binding site for a specific TF in region [-250,0] upstream the binding site of the target TF, while the sequences of the second class have not these potential binding sites. Hence, the goal of this module is to identify, for each PWM of the library, a specific region of the sequences in which the scores of this PWM are higher in one class than in the other one.

Sequences are first segmented in bins of the same size. We used 13 bins in the following experiments. The number of bins impacts the precision of the approach but also the computing time of the analysis. For each PWM, TFscope scans all sequences with FIMO [18], and the best score achieved on each bin of each sequence is stored. Then, TFscope searches the region of consecutive bins for which the PWM gets the most different scores depending on the class of the sequences. A lattice structure is used for this exploration (see Figure 1 and details in section Materials and Methods). For each PWM of the library, TFscope-CF selects the region that shows the highest differences and returns a variable corresponding to this PWM and region. As for TFscope-NE, the core-motif is masked before running the analysis.

### Analysis of the cellular specificities of 272 ChIP-seq pairs

We first sought to apply TFscope to identify binding sites differences of TFs in different cell types using a selection of 272 pairs of ChIP-seq experiments downloaded from the GTRD database [25]. To minimize the effects linked to technical issues or indirect binding, data were filtered using the UniBind p-value score [16]. In UniBind the authors studied the distance between the ChIP-seq peaks and the position of the most likely binding site (inferred with the PFM associated with the TF studied). They showed that this binding site is sometimes far from the ChIP-seq peak, and that the peak may be a false positive. Using a dedicated method named ChIP-eat, the authors were able to determine genomic boundaries inside which the binding sites are likely true positives, and they provide a p-value measuring peak enrichment in these boundaries. We used this p-value to remove ChIP-seq experiments that could be affected by technical issues and indirect binding. Moreover, we only selected for this analysis pairs of experiments that show strong binding site differences according to Jaccard’s distance (see Materials and method). The 272 pairs were chosen to provide a wide view of the ChIP-seq data in GTRD, *i*.*e*. pairs that were too close to another pair already selected were discarded (see the pair selection procedure in Materials and method).

### TFscope learns both discriminative and informative core motifs

We first assessed the TFscope ability to identify core motif differences in the ChIP-seq experiments pairs. In this analysis, we only used the score function of the learned PWM (Expression (2)) to discriminate the two cell-types. For comparison, we also used the score of the original PWM on this problem. Accuracies were measured by AUROC on an independent set of sequences (see Figure 2A). If several versions of the original PWM were available, we used the version that provides the best AUROC. As we can see, the new PWM outperforms the original PWM most of the time. Moreover, we can also observe that for some of the 272 experiment pairs, the core motif itself is sufficient to differentiate the two cell-types with high accuracy. As already discussed, the discriminative PWM is different from the original PWM, as it specifically models the differences while removing the features common to the two classes. The “mirror-logo” representation summarizes these differences and shows which features are associated with which cell-type. For example, the Figure 2B (up) shows the original CEBPA PWM provided by JASPAR, while the middle figure shows the mirror logo of the discriminative PWM learned by TFscope for discriminating CEBPM binding sites between the SKH1 and U937 cell-types. One can see here that the canonical CEBPA motif is more often associated with ChIP-seq peaks collected in U937 than in SKH1. Although the SKH1 sequences also bear a very similar motif (recall that the motif is present at the center of the sequences for both conditions) the mirror logo indicates for example that the T nucleotides at positions 3 and 4 are more often missing in the SKH1 sequences than in the U937 sequences. Similarly, among the small differences that may exist between the motifs in the two conditions, it seems that the SKH1 sequences often have a C at position 5.

We next sought to compare these results to those obtained with another method that learns discriminative PWMs. We used the DAMO approach for this comparison, as it is one of the rare methods that do not rely on PPM to learn a PWM. Recall that DAMO, as the other classical approaches to learn PWMs, has not been designed to address exactly the same problem as our. Indeed, DAMO usually takes in input sequences that are not aligned and that are much longer than the target PWM. Nevertheless, it can also be used on our simpler problem. However, as illustrated on Figure 2C, it does not achieve the same accuracy as TFscope on this problem, which was somewhat expected as the logistic classifier used by TFscope theoretically returns the most discriminative PWM.

Another striking fact when we compare the discriminative motifs learned by DAMO to those of TFscope is that the DAMO motifs appear much more complex, with a lot of positions without clear preferences. On the contrary, thanks to the LASSO penalty used for learning, the TFscope motifs are easier to interpret, with many positions set to zero (compare the two examples provided on Figure 2B). This aspect was assessed systematically on the 272 experiments using a score function based on the Gini coefficient for measuring motif simplicity (see Material and methods). As illustrated on Figure 2D, TFscope motifs have higher Gini coefficient, and are thus simpler and easier to interpret than their DAMO counterpart.

Finally, we observed that increasing the size of the PWM until 4 nucleotides on both sides still improves the AUROC of the DM model (see Supp. Fig. 1). So, in the following, this model (denoted as DM+8) is used in the TFscope model and the experiments.

### Position of co-factors helps for predicting cell-specificity

We next sought to investigate the information gained by the position of the binding sites of potential co-factors for cell-type prediction. We used for this a simplification of the model of Expression (1), which only uses the core motif and the co-factor variables for the prediction— *i*.*e*. the NE_*i*_ variables capturing the nucleotidic environment were removed from the model. The accuracy of this model was compared to that of a similar model that also uses the score of potential co-factors, but without integrating the information of position. This model, which strongly resembles the TFcoop approach we previously proposed [47], simply uses the best score achieved by the different PWMs in the whole sequence. Hence, the predictive variables of this model are the best scores achieved at any position of the sequence, while in Expression (1) TFscope uses the best score achieved in a specific region identified as the most informative for each cofactor. While the two models have exactly the same number of parameters (*i*.*e*. the number of PWMs in the PWM library), the variables of TFscope greatly increase the accuracy of the approach (Figure 3), illustrating the fact that position of co-factors relative to the considered TFBS also carry important information. Note that, as we will see in the following, TFscope provides a graphical representation of all identified co-factors, and position information can be easily retrieved.

**Figure 3.**
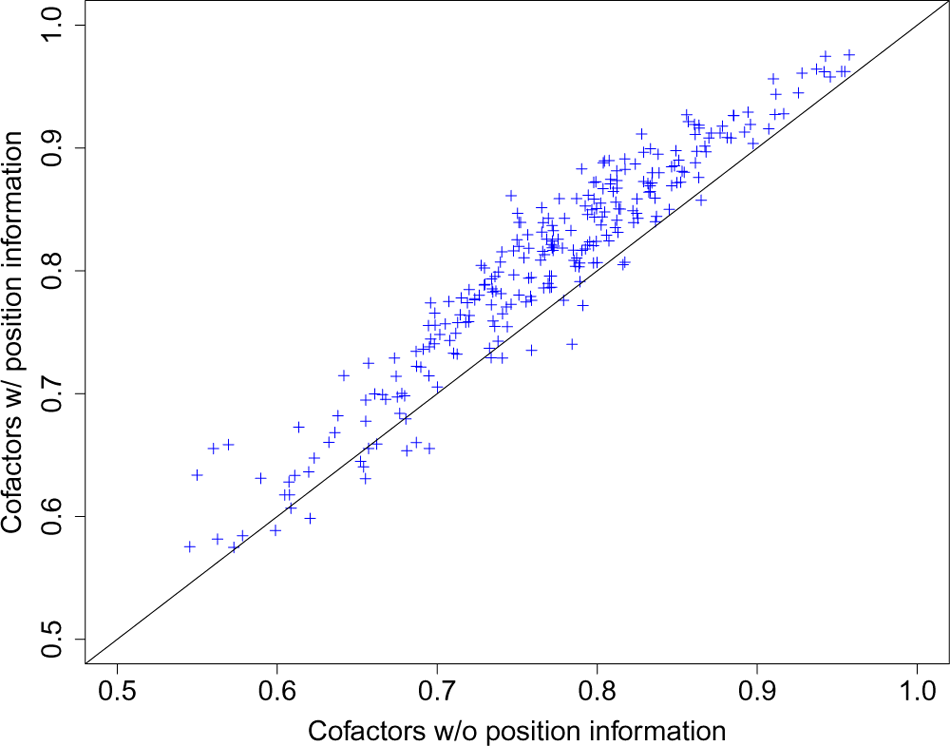
Position of co-factors helps for predicting cell-specificity. AUROCs achieved by TFscope models that only use the score of co-factors for the predictions. On the x-axis, the score of a co-factor on a sequence *s* is the best achieved at any position of the sequence, *i*.*e*. the position of the co-factor is not considered in the score computation. On the y-axis, the score of each co-factor is only computed on a specific region identified by the TFscope-CF module to be the most informative for this co-factor.

### TFscope assesses the relative importance of each genomic feature

We next ran TFscope with the full model of Expression (1) on the 272 pairs of experiments and compared its accuracy (AUROC) to that of different alternatives: the original PWM only, the discriminative PWM only, and three incomplete TFscope models that only use two of the three kinds of genomic information. These incomplete models were obtained by taking the full TFscope model trained with all variables, and by setting to zero either the variable DM (model TFscope w/o core motif information), or the NE_*i*_ variables (TFscope w/o nucleotidic environment information), or the CF_*j*_ variables (TFscope w/o co-factor information). Figure 4A reports the accuracy achieved by all these models. As we can see, the full TFscope model successfully integrates the three kinds of genomic information and outperforms the alternative models. We can also observe that the accuracy is often good, with a median AUROC above 80%. Moreover, there is a strong link between the accuracy of the approach and the Jaccard distance between the ChIP-seq peaks in the two cell types (Pearson *r*=0.51; see Figure 4B), *i*.*e*. experiments with low proportion of ChIP-seq peaks shared by the two cell types often get good accuracy (remember that these peaks are removed before the analyses). In other words, when the two ChIP-seq experiments are really different, TFscope accurately predicts these differences. Figure 4B also illustrates the fact that, for most analyses, the Jaccard distance is high (so the Jaccard index is low, see Supp. Fig. 2). This means that the number of common peaks is small in proportion, hence removing these peaks makes sense for our analyses.

**Figure 4.**
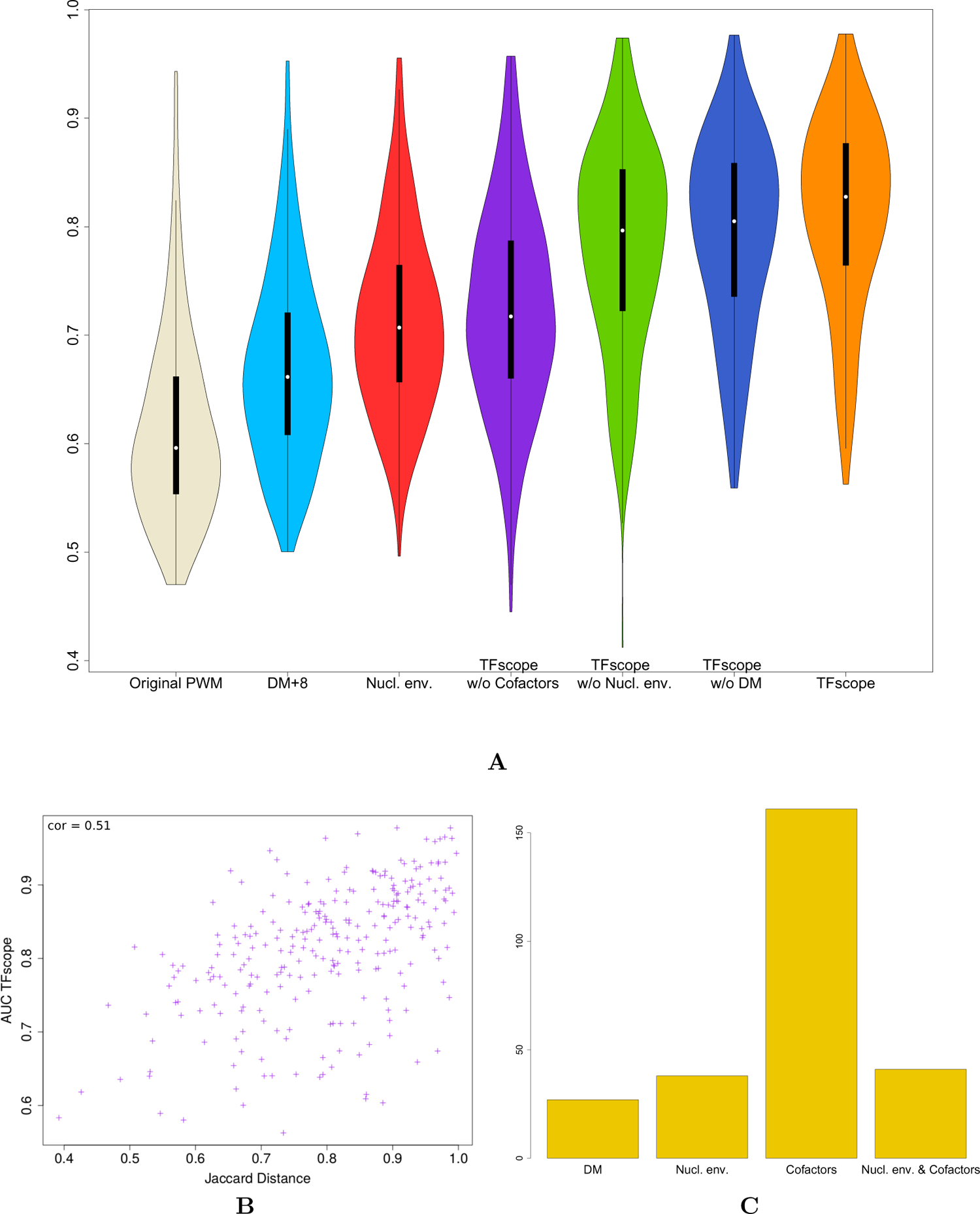
Accuracy achieved by TFscope for discriminating binding sites of different cell types. **A** Distribution of AUROCs achieved by TFscope and several alternative models for discriminating binding sites of one TF in two different cell types. **B** Link between TFscope accuracy and the similarity of ChIP-seq peaks in the two cell types. ChIP-seq experiments that have a high proportion of peaks in common have low Jaccard distance (Jaccard distance = 1 - Jaccard index). **C** Distribution of TFscope models according to what are the most discriminative features: the discriminative motif (DM), the nucleotidic environment, the co-factors, or the nucleotidic environment + co-factors.

These good performances legitimate the use of TFscope to investigate the relative importance of each kind of genomic information in the different comparisons. For this, in addition to the logo of the discriminative PWM, TFscope outputs a radar plot that summarizes the accuracy of the different models and alternatives, and a location plot that summarizes the position of the most important variables of the model (see Material and methods). For example, Figure 5B reports the radar plot obtained when analyzing the binding differences of TF JUND between liver and lung carcinoma. For this experiment, the core motif is clearly the most discriminant information (Figure 5A), since removing this information lead to the largest drop of AUROC. Besides, peaks detected in lung harbor additional AP-1 motifs around the core motif (Figure 5C). JunD belongs to the AP-1 family of dimeric TFs, which associate members of the Jun (c-Jun, JunB and JunD) and Fos (c-Fos, FosB, Fra-1/Fosl1 and Fra-2/Fosl2) families. In contrast to the Jun family members, which can homodimerize, the Fos family members must heterodimerize with one of the Jun proteins to bind DNA. Importantly, Fos:Jun heterodimers have a stronger affinity for DNA than the Jun:Jun homodimers [7]. According to various expression data listed in the EBI Expression Atlas (https://www.ebi.ac.uk/gxa/home), Fos TFs are less expressed in liver than in lung. Thus, JunD binding preferences observed in liver *vs*. lung might merely be explained by the expression of Fos TFs: because the probability to form Fos:Jun heterodimers is greater in lung than in liver, JunD will bind DNA with a higher affinity in lung than in liver. For comparison, the discriminative motif and the radar plot of the CTCF experiment between CD20 and RH4 are on Figure 5D-E. Here, the most discriminative information seems to be the nucleotidic environment. The location plot provides additional information (Figure 5F). We can see that CD20 favors A/T rich environment in the vicinity of the binding motif (∼ +/ − 100bp around the motif), and C/G nucleotides in the larger surrounding region (+/ − 500bp). On the contrary, RH4 prefers a nucleotide environment rich in TG and CA dinucleotides. All results obtained on the 272 experiments are available on https://gite.lirmm.fr/rromero/tfscope/-/tree/main/results.

**Figure 5.**
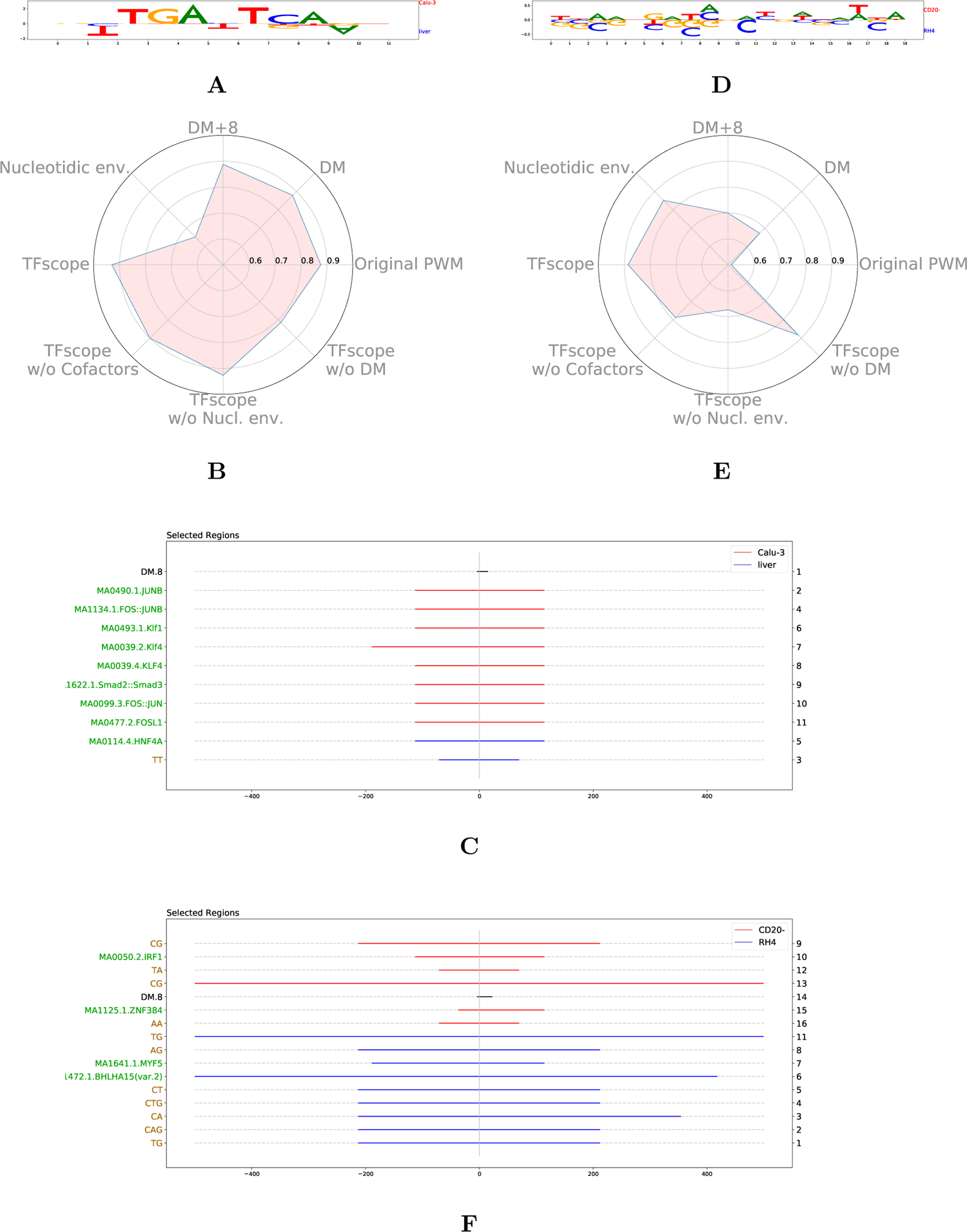
Core motif, nucleotidic environment and co-factors together determine cell specificity. **A-B-C** Discriminative PWM, radar plot and location of the most important variables in the JUND comparison between liver and lung carcinoma. **D-E-F** Discriminative PWM, radar plot and location of the most important variables in the CTCF comparison between B lymphocite and rhabdomyosarcoma. Radar plots (B & E) summarize the AUROC achieved by TFscope and several alternative models. Location plots (C & F) provide the identity and location of the most important variables (black: DM; green: co-factors; brown: nucleotidic environment). The numbers on the right hand indicate the ranking of the variables, from the most important (rank 1) to the least important. The color of segments indicates the cell-type associated with each variable.

Finally, in an attempt to provide a broad picture of the genomic strategies involved in the control of binding differences between cell types, we ran a K-means clustering on the importance profiles inferred by TFscope. More precisely, all 272 experiments were described by a vector of length 3 obtained by subtracting the AUROC of TFscope w/o DM, TFscope w/o NE and TFscope w/o CF from that of the full TFscope model. Each experiment is then represented by three values representing the three AUROC losses associated with the three kinds of information. We reasoned that a maximum of 7 broad classes can be expected from these data: three classes with a single information clearly higher than the two others, three classes with two more-important information, and one class with approximately equal importance of the three information. However, by visually inspecting the results of several K-means clustering, we end up with a total of only 4 classes: the three-information class, and the DM+NE and DM+CF classes seem absent from the 272 models. Figure 4C reports the distribution of the 272 models in these 4 classes, highlighting the fact that the co-factors are by far the most common mechanism involved in the binding differences between cell types. The target motif itself appears to be the more discriminative feature in 10.6% of the 272 experiments, and the nucleotidic environment around the binding site in 14% cases.

### Analysis of the binding differences induced by a specific treatment

We next sought to use TFscope to analyze the binding differences observed between two ChIP-seq experiments targeting the same cell type but with two different treatments. 79 ChIP-seq pairs were selected (see Material & Methods) and analyzed. As for the cell type comparisons, all results obtained on the 79 experiments are available on https://gite.lirmm.fr/rromero/tfscope/-/tree/main/results. We got globally similar results than for the cell type experiments (see the plot of accuracy in Supp. Fig.3A), although the Jaccard distance between treatments is often smaller than between cell types (Supp. Fig.3B), *i*.*e*. two treatments often show more similar binding sites than two cell types. However, for several experiments there is a clear difference in the binding sites and TFscope indentifies interesting features.

For instance, TFscope confirms the cross-talk between GR signaling and NF-*κ*B reported in [23] and proposes additional features. Specifically, analyzing NR3C1 ChIP-seq upon Dexamethasone (Dex) and Dex+TNF treatments with TFscope reveals that the main features distinguishing the binding sites in these two conditions are cooperating TFs (Figure 6A). While motifs of NFI-related TFs are enriched in NR3C1 peaks upon Dex treatment alone, as observed in [23], motifs of NF-*κ*B-related TFs are enriched in NR3C1 peaks upon Dex+TNF (Figure 6B).

**Figure 6.**
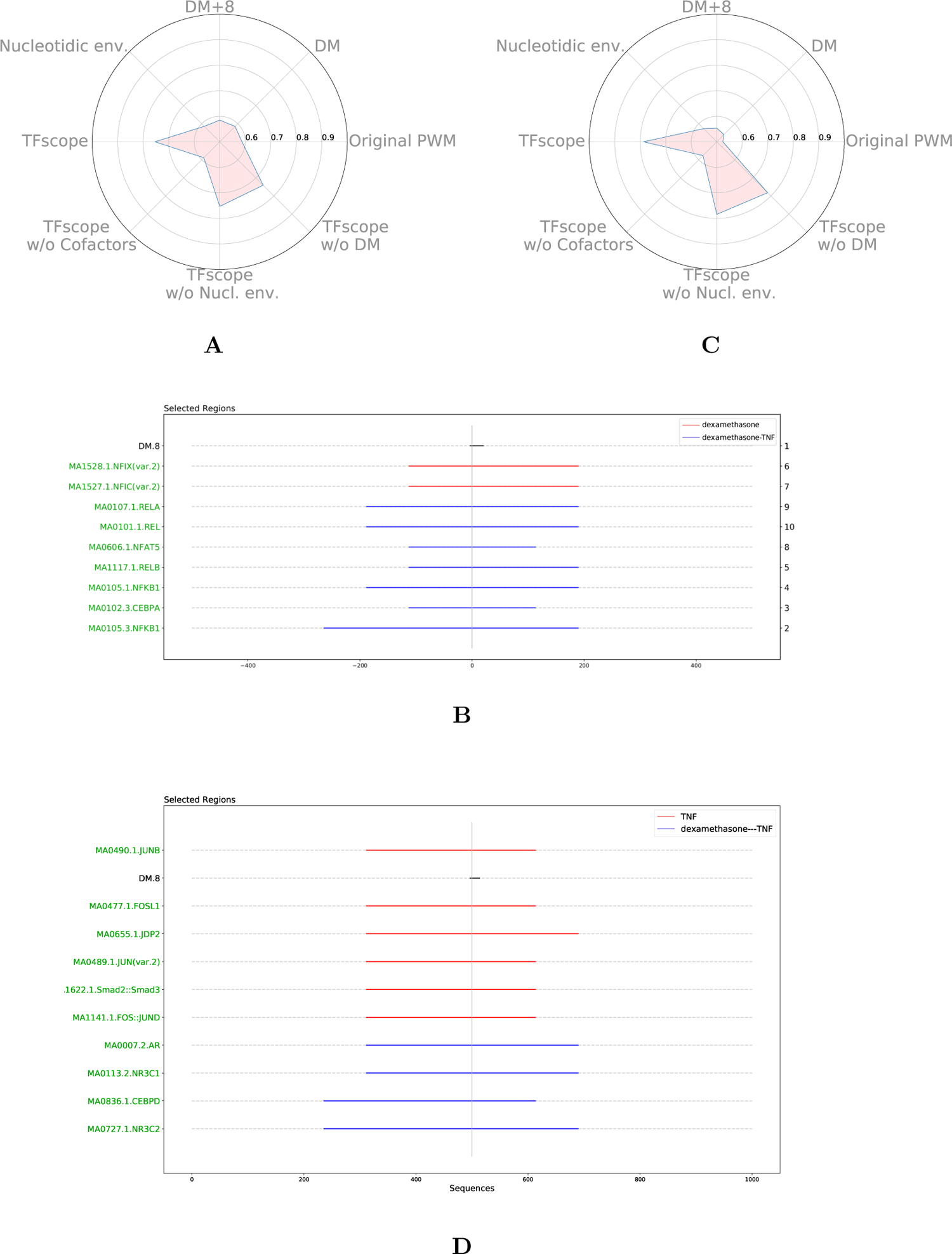
Analysis of the binding differences induced by a specific treatment. **A-B** Radar plot and location plot of the most important variables in the NR3C1 comparison between Dex and Dex+TNF treatments. **C-D** Radar plot and location plot of the most important variables in the RELA comparison between TNF and Dex+TNF treatments.

Similarly, cooperating TFs appear as the main features distinguishing RELA ChIP-seq peaks upon TNF and Dex+TNF treatments (Figure 6C). TFscope confirms that, in the presence of Dex, RELA peaks are associated with steroid receptor TFs (NR3C1, NR3C2 and AR) but it also suggests that GR signaling abolishes cooperation with AP-1 related TFs observed preferentially in RELA peaks in pro-inflammatory conditions (TNF alone) (Figure 6D).

### Analysis of the binding differences of paralogous TFs

We showed in a previous work [8] that the binding of two paralogous TFs, namely FOSL1 and FOSL2 (also called as FRA1 and FRA2), can be distinguished primarily by the scores of their motif: FOSL2 preferentially binds sequences harboring high scores for the canonical AP-1 motif, while FOSL1 binds sequences with some degenerate positions (lower scores). We then thought to use TFscope to distinguish FOSL1 from FOSL2 binding on the same dataset. TFscope-DM is indeed sufficient to classify the two peak classes (Figure 7A) and the typical AP-1 motif is more frequently found in FOSL2 peaks (Figure 7B), confirming our previous results obtained with another approach [47]. Moreover, the discriminative motif also brings new information. For example, FOSL1 favors nucleotides that are inverse from those of the canonical motif in positions 2 and 10.

**Figure 7.**
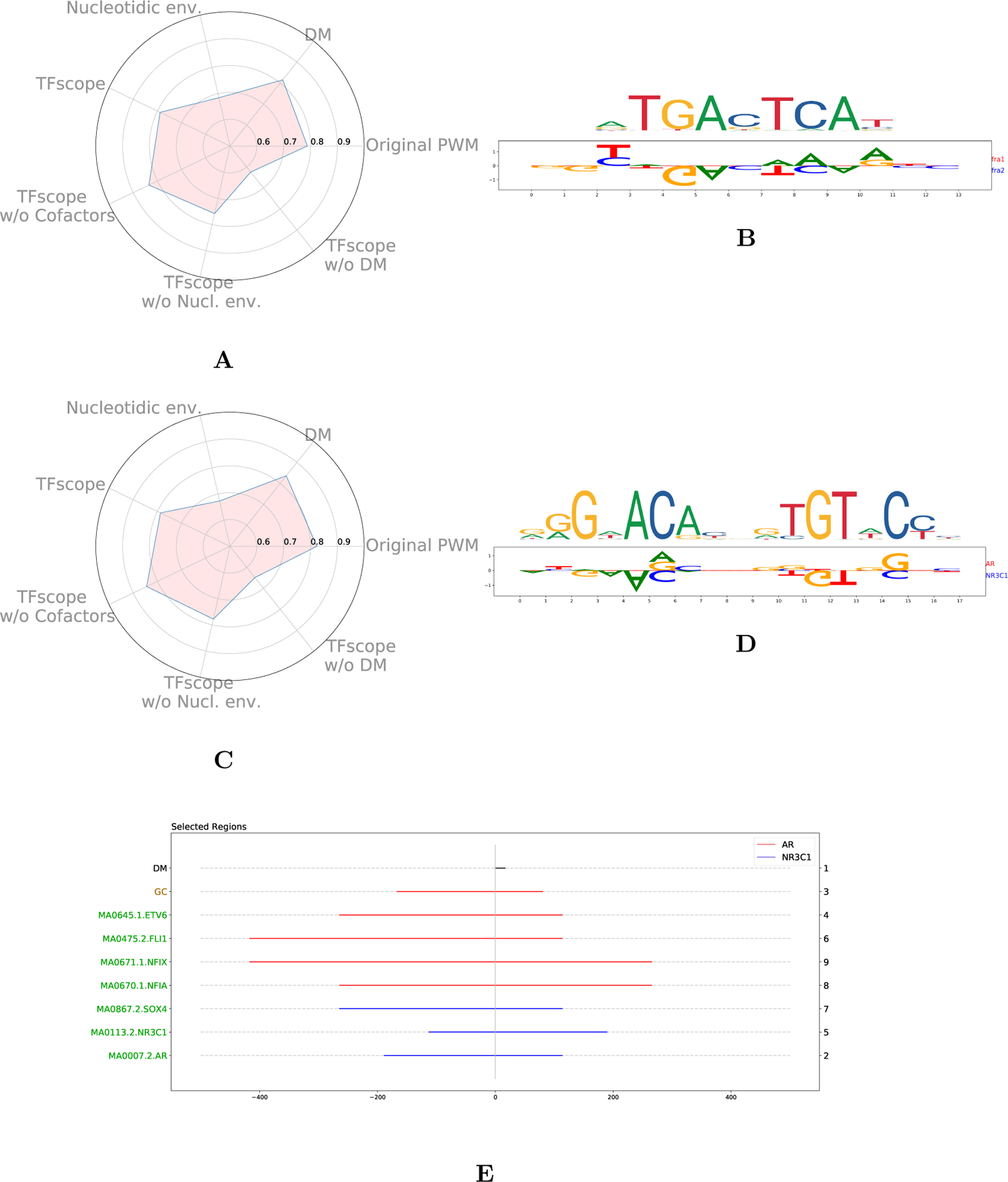
Analysis of the binding differences of paralogous TFs. **A** Radar plot of the most important variables discriminating FOSL1 and FOLS2 binding sites. **B** JASPAR FOSL1 motif (up) and TFscope-DM motif discriminating FOSL1 and FOLS2 binding sites (down) **C** Radar plot of the most important variables discriminating AR and GR binding sites. **D** JASPAR AR motif (up) and TFscope-DM motif discriminating AR and GR (NR3C1) binding sites (down). **E** Location plot of variables discriminating AR and GR binding sites.

To confirm the applicability of TFscope in this sort of classification task, we considered another pair of paralogous TFs, NR3C1 and AR, and ChIP-seq data collected in MCF-7 cells [41]. As shown in Figure 7C, TFscope is able to accurately distinguish NR3C1 from AR ChIP peaks, and TFscope-DM shows that the main differences lie in the core motif itself. The output of TFscope-DM reveals that dinucleotides AC at position 4 and GT at position 11 in the canonical NR3C1/AR motif are more frequent for NR3C1 than for AR (Figure 7D). Moreover, AR ChIP-seq peaks appear more GC-rich than NR3C1 peaks (Figure 7E). These results are in full agreement with that obtained by Kulik *et al*., who compared AR and GR binding preferences in U2OS cells [29]. Together these results illustrate the possibility to use TFscope to distinguish the binding of paralogous TFs.

## Discussion

We proposed here a new machine learning approach to identify the DNA features that can explain the binding preferences of a TF in two settings: two cell types, two conditions, or two paralogous TFs. Our approach uses three modules that identify three kinds of DNA features related to TF binding. The first one is a new method to learn a discriminative PWM. Among the numerous approaches already proposed to learn a PWM, it is important to note that most of them actually learn a PPM which is then converted into a PWM with a simple log ratio formula (see for example reference [49]). The problem with this procedure is that it potentially impedes the accuracy of the PWM. Indeed, PPMs being probabilistic models, they are subject to strong constraints (notably, the sum of a PPM column must be equal to one) which inevitably also constrains the weights of the PWM. For example, the log-ratio operation cannot produce a PWM in which one of the columns has all but one weight equal to zero (the log ratio gives zero when the probability of the nucleotide at this position equals the probability of the nucleotide in the background; but if it is the case for 3 nucleotides it is also necessarily the case for the 4th nucleotide). Interested readers can refer to the work of Ruan and Stormo [38] for more arguments about the limits of PPMs for PWM learning. Our approach based on logistic regression avoids this problem and has moreover the advantage of allowing us to include a LASSO penalty to get simpler and more readable PWMs. Another important difference with previous approaches is that in TFscope the discriminative PWM is only used to discriminate the two classes, but not to scan the sequences. Hence TFscope needs two PWMs: the JASPAR PWM is used to scan the sequences and identify the binding sites in both classes, and the discriminative PWM is used to score the binding sites and differentiate the two classes.

In addition to the specificity of the core motif that is captured by our discriminative PWM, the two other modules extract DNA features related to the nucleotidic environment around the TFBS, and the presence and position of every potential co-factor. Then, a learning algorithm is run to both train a model and select the most discriminative features at the same time. Hence, contrary to the CNNs based methods that have been recently proposed, our approach completely controls the predictive features used by the model. This allows us to easily assess the global importance of each feature, by measuring the loss of accuracy induced by its withdrawal, something very challenging to do with CNN approaches.

Our results on different TFs and different cell types show that co-factors are often the most important determinant associated with the cell-specific binding sites, and that their position relative to the TFBS considered is key. However, for several experiments such as CTCF in CD20 *vs*. RH4 the large nucleotidic environment around the binding sites also explains a part of the observed differences. For some other experiments such as JUND in lung *vs*. liver the main differences lie directly in specific nucleotides of the binding site. When comparing two treatments the picture is globally the same, while for paralogous TFs the main differences are associated with the core motifs themselves in our experiments. In this latter case, although the binding motifs globally show very similar PWMs for both TFs, subtle differences at specific positions actually explain most of the binding differences.

Our approach could be improved in different ways. Notably, one drawback that can sometimes hamper a straight interpretation of the TFscope results is the correlation between predictive variables. Scores of TF motifs especially may be highly correlated, as several TFs often share very similar motifs. Hence, although the PWM highlighted by TFscope is the one that shows the highest link with the predicted signal, other PWMs could also have a high correlation, and thus other TFs are potential co-factors. We therefore encourage users to refer to PWM clusters as defined for example in the RSAT-matrix clustering [10]. Similarly, there are sometimes correlations between the nucleotidic composition captured by the TFscope-NE module and the co-factor motifs identified by TFscope-CF. Here again, the linear model and the LASSO penalty ensure that the variables selected by TFscope are those with the strongest link with the predicted signal. Nevertheless, it is important to keep in mind that other variables may actually be involved in the studied process. We are thus working on a way to identify and present all alternative variables in a friendly interface. Another improvement would be to integrate additional DNA features into our model. Specifically, the number of repeats of a given PWM could be an interesting variable for discriminating two ChIP-seq experiments. Such information is not directly accounted for in the current model but could potentially explain binding differences in some experiments.

## Material and Methods

### Sequence extraction and alignment

TFscope takes in input two sets of ChIP-seq peaks provided as BED files. First, all peaks common to the two files are removed. This is done with the BED tools using

~~~
bedtools window -v -w 500 -a class0.bed -b class1.bed > class0_no_overlap.bed
bedtools window -v -w 500 -a class1.bed -b class0.bed > class1_no_overlap.bed
~~~

Then the sequences corresponding to each peak are extracted and aligned on the most likely occurrence of the TFBS. We use for this the PWM associated with the target TF in JASPAR 2020 [15]. FIMO is used to parse the sequences with the command

~~~
fimo --thresh 0.001 --max-strand --text --bfile background_fimo.txt
 PPM_jaspar2020.meme fasta.fa > occurrences.dat
~~~

The best occurrence of the motif around the ChIP-seq peak (in a limit of 500kb) is identified and used as an anchor point around which the 1Kb sequences are centered (see Figure 1). Sequences for which no occurrence of the motif is found around the peak summit are discarded. Finally, the number of sequences of the two classes are rebalanced, *i*.*e*. some sequences of the larger class are randomly selected and removed, in order to get two classes with an equal number of sequences.

If several versions of the PWM are available in JASPAR, we used the PWM that is the most discriminative for the problem at hand. Namely, for each PWM, the best occurrence of the motif is identified on each sequence, and these scores are used to discriminate the two classes (this corresponds to the AUROC of the original PWM in the radar plots). The PWM version with the highest AUROC is used for the rest of the analysis.

The formatted data used in the experiments are available in the dedicated gite repository: https://gite.lirmm.fr/rromero/tfscope.

### TFscope-DM

This module takes as input the *K*-length sub-sequence corresponding to the most likely occurrence of the motif in each sequence (*K* being the size of the PWM). Each sub-sequence *s* is one-hot encoded in a *K ×* 4 matrix **s**. Then, a logistic model with *K ×* 4 parameters (see Expression 2) is learned to discriminate the two classes of sub-sequences. The parameters of the model are estimated by maximum likelihood, with a LASSO penalization [46] to favor simple and easy-to-interpret models. This is done with library glmnet on python 3.

### TFscope-NE

This module takes in input the 1Kb sequences centered on the most likely TFBS (cf. Sequence extraction and alignment). The *K*-length sub-sequence corresponding to the TFBS is masked (replaced by *K* N nucleotides) to avoid capturing information related to the core-motif. Then the TFscope-NE module constructs new variables defined by a pair (kmer,region) such that the frequency of the identified k-mer in the associated region is, on average, different between the two classes. We used for this a slight modification of the DExTER method [32]: rather than searching for variables that are correlated with an expression signal, TFscope-NE extracts variables that can discriminate the two classes, as measured by the AUROC. The rest of the procedure is exactly the same as that used in DExTER (see ref. [32] for details). Sequences are first segmented into different bins. We used 7 bins in the experiments. TFscope-NE starts with 2-mer (dinucleotides) and, for each 2-mer, identifies the region of consecutive bins for which the 2-mer frequency in the region is the most discriminant. Once the best region has been identified for a 2-mer, TFscope-NE attempts to iteratively extend this 2-mer for identifying longer k-mers (up to 4-mers). At the end of the process, a set of variables corresponding to the frequency of the identified k-mers in the identified regions is returned for each sequence.

### TFscope-CF

As TFscope-NE, this module takes in input the 1Kb sequences centered on the most likely TFBS (this TFBS is also masked to avoid capturing information related to the core-motif). This module constructs variables defined by a pair (PWM,region) such that the score of the PWM in the identified region is, on average, different between the two classes. For this, sequences are first segmented in bins of the same size. We used 13 bins in the experiments. The number of bins impacts the precision of the approach but also the computing time of the analysis. For each PWM, TFscope scans all sequences with FIMO, and the best score achieved in each bin of each sequence is stored. Then, TFscope uses a lattice structure (see Figure 1) to compute the best score achieved in any region made up of consecutive bins. Each node of the lattice is associated with a specific region: the top of the lattice represents the whole sequence, while the lowest nodes represent the different bins. Once the best score achieved in every bin has been computed, the best score achieved in any node of the lattice can be easily deduced with a max() operation on its two children nodes. For example, the lattice of Figure 1 corresponds to a sequence for which the best score is obtained in the first bin ([-500;-300]). For each PWM, a lattice like this one is computed for every sequence. Then, TFscope identifies the node (region) such that the scores associated with this node in the different lattices provide the highest AUROC for discriminating the two classes.

### Selection of 272 ChIP-seq pairs targeting the same TF in two different cell types

272 pairs of experiments targeting a common TF, with the same treatment, in two different cell-types were selected from the GTRD and UniBind databases. ChIP-seq data were downloaded from GTRD http://gtrd20-06.biouml.org/downloads/20.06/bigBeds/hg38/ChIP-seq/Clusters_by_TF_and_Peakcaller/MACS2/. Only experiments associated with a UniBind p-value below 1% were considered, which represents a total of 2815 ChIP-seq data. This data can be arranged in a total of 6553 pairs targeting a common TF, with the same treatment, in two different cell-types. We chose to select only the pairs that show highly different peaks for the analyses. This was measured with the Jaccard’s distance. Let A and B be two sets of ChIP-seq peaks on the genome, the Jaccard’s distance *D*_*J*_ is defined from the Jaccard index by:

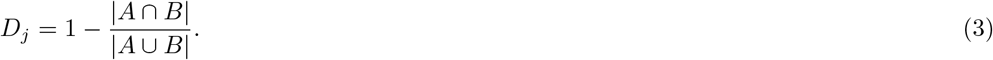

Peak intersections and unions were computed with Bedtools window and merge, respectively:

~~~
bedtools window -w 500 -a class0.bed -b class1.bed > intersection.bed
bedtools merge -d 500 -i intersection.bed
~~~

For a given TF and treatment, several pairs with different cell-types are often possible. In order to select only a subset of these pairs, we ran a hierarchical clustering of all the data targeting the same TF with the same treatment. The clustering was done using the Jaccard’s distance and the complete-linkage agglomeration strategy. We then selected one pair of experiments for each internal node of the tree (the two experiments with the highest number of peaks were selected). Hence, if the tree has *N* leaves (corresponding to the *N* ChIP-seq experiments targeting the same TF and treatment) the number of pairs is exactly *N*. In this way, we end up with a total of 425 ChIP-seq pairs, which were reduced to 368 pairs by selecting only pairs with at least 1000 specific peaks in each cell-type. Among these pairs, more than 100 actually involved CTCF. We chose to keep only 7 CTCF pairs (which were chosen as the pairs with the largest Jaccard distance), and we end up with a final set of 272 pairs of experiments.

### Measure of PWM simplicity

Information content derived from information theory is often used to measure the conservation of specific nucleotides at specific positions of a motif. This measure is however based on probability distribution and is thus restricted to PPMs: it does not extend to the PWM general case. Hence, we propose to use the Gini coefficient to measure PWM simplicity.

The Lorenz curve is a graphical representation of the income distribution between individuals in econometrics. It is obtained by ordering the individuals in the order of their income, and by calculating the cumulative part of income in function of the cumulative part of individuals (see Supp. Figure 4). In the case of an equal distribution between all the individuals, the curve follows the line *y* = *x*. Otherwise, it is found under this line. The surface *A* is the area between the Lorenz curve and the line of perfect equality of distribution. The surface *B* is the area between the Lorenz curve and the perfect inequality curve (all the income belongs to a single individual). The Gini coefficient is defined by *A/*(*A* + *B*). It is equal to 1 if all the incomes belong to a single individual and equal to 0 if the incomes are equally distributed.

For PWMs, we compute the Gini coefficient on the set of PWM weights. More precisely, we gather the 4 *× K* weights of the PWM (all positions combined), order them in ascending order of their absolute value, and compute the Lorenz curve and the Gini coefficient associated with this set of weights. A small Gini coefficient implies an equal distribution of the PWM weights: the information is dispersed on many elements of the PWM. On the contrary, a large Gini coefficient (close to 1) indicates that some weights of the PWM gather all the information and that many weights are equal or close to 0. So we can see it as a measure of the interpretability of the models, where a model with a large Gini coefficient will be simpler than a model with a Gini coefficient close to 0. Importantly, the Gini coefficient does not depend on the scale of the PWM weights but only on their relative importance. Hence, it can be used to compare PWMs obtained from different methods.

### Measure of variable importance

We devised an *ad hoc* procedure based on LASSO penalty and model error for measuring the individual importance of the different variables of a model. Given a penalization constraint *λ*, the LASSO procedure searches the model parameters that minimize the prediction error subject to the constraint. In practice, a grid of constraints of decreasing values is initialized, and a model is learned for each value. The result is a series of models with an increasing number of parameters. To identify the most important variables of a model in a given condition, we took the model with 10 parameters and estimated the importance of each of the 10 variables in the following way. Given a variable *X*, its importance was estimated by the AUROC difference between the complete model and the model obtained by setting *β*_*X*_ to 0.

### Selection of 79 ChIP-seq pairs targeting the same TF with two different treatments

To compare binding preferences upon different treatments, we removed experiments associated with the ‘no-condition’ term in our GTRD/UniBind joint list. We sorted the remaining 1, 354 experiments according to their GTRD IDs in order to consider experiments from the same publication/study. We further removed time-course experiments and selected 100 pairs of possible comparisons. Then, the same procedure as the one used for the selection of the 272 ChIP-seq pairs was applied. This gives a total of 79 pairs of ChIP-seq experiments.

## Acknowledgements

We thank Marius Gheorghe and Anthony Mathelier for their helpful comments and assistance with Unibind data.

## References

[1] Ariel Afek, Hila Cohen, Shiran Barber-Zucker, Raluca Gordân, and David B. Lukatsky. Non-consensus Protein Binding to Repetitive DNA Sequence Elements Significantly Affects Eukaryotic Genomes. PLOS Computational Biology, 11(8):e1004429, August 2015. Publisher: Public Library of Science.

[2] Vikram Agarwal and Jay Shendure. Predicting mRNA Abundance Directly from Genomic Sequence Using Deep Convolutional Neural Networks. Cell Reports, 31(7):107663, May 2020.

[3] David N. Arnosti and Meghana M. Kulkarni. Transcriptional enhancers: Intelligent enhanceosomes or flexible billboards? Journal of Cellular Biochemistry, 94(5):890–898, April 2005.

[4] Žiga Avsec, Melanie Weilert, Avanti Shrikumar, Sabrina Krueger, Amr Alexandari, Khyati Dalal, Robin Fropf, Charles McAnany, Julien Gagneur, Anshul Kundaje, and Julia Zeitlinger. Base-resolution models of transcription-factor binding reveal soft motif syntax. Nature Genetics, 53(3):354–366, March 2021. Bandiera abtest: a Cg type: Nature Research Journals Number: 3 Primary_atype: Research Publisher: Nature Publishing Group Subject term: Chromatin immunoprecipitation;Computational biology and bioinformatics;Genomics Subject term id: chromatin-immunoprecipitation;computational-biology-and-bioinformatics;genomics.

[5] Timothy L Bailey. STREME: accurate and versatile sequence motif discovery. Bioinformatics, (btab203), March 2021.

[6] Timothy L. Bailey and Philip Machanick. Inferring direct DNA binding from ChIP-seq. Nucleic Acids Research, 40(17):e128, September 2012.

[7] Fabienne Bejjani, Emilie Evanno, Kazem Zibara, Marc Piechaczyk, and Isabelle Jariel-Encontre. The AP-1 transcriptional complex: Local switch or remote command? Biochimica et Biophysica Acta (BBA)-Reviews on Cancer, 1872(1):11–23, 2019. Publisher: Elsevier.

[8] Fabienne Bejjani, Claire Tolza, Mathias Boulanger, Damien Downes, Raphaël Romero, Muhammad Ahmad Maqbool, Amal Zine El Aabidine, Jean-Christophe Andrau, Sophie Lebre, Laurent Brehelin, Hughes Parrinello, Marine Rohmer, Tony Kaoma, Laurent Vallar, Jim R Hughes, Kazem Zibara, Charles-Henri Lecellier, Marc Piechaczyk, and Isabelle Jariel-Encontre. Fra-1 regulates its target genes via binding to remote enhancers without exerting major control on chromatin architecture in triple negative breast cancers. Nucleic acids research, 49(5):2488–2508, 2021. Publisher: Oxford University Press.

[9] Milagros Castellanos, Nivin Mothi, and Victor Muñoz. Eukaryotic transcription factors can track and control their target genes using DNA antennas. Nature Communications, 11, January 2020.

[10] Jaime Abraham Castro-Mondragon, Sébastien Jaeger, Denis Thieffry, Morgane Thomas-Chollier, and Jacques van Helden. RSAT matrix-clustering: dynamic exploration and redundancy reduction of transcription factor binding motif collections. Nucleic Acids Research, 45(13):e119, July 2017.

[11] Hemangi G. Chaudhari and Barak A. Cohen. Local sequence features that influence AP-1 cis-regulatory activity. Genome Research, 28(2):171–181, February 2018.

[12] Iris Dror, Tamar Golan, Carmit Levy, Remo Rohs, and Yael Mandel-Gutfreund. A widespread role of the motif environment in transcription factor binding across diverse protein families. Genome Research, 25(9):1268–1280, January 2015.

[13] ENCODE Project Consortium. The ENCODE (ENCyclopedia Of DNA Elements) Project. Science (New York, N.Y.), 306(5696):636–640, October 2004.

[14] Jason Ernst and Manolis Kellis. Interplay between chromatin state, regulator binding, and regulatory motifs in six human cell types. Genome Research, 23(7):1142–1154, July 2013.

[15] Oriol Fornes, Jaime A. Castro-Mondragon, Aziz Khan, Robin van der Lee, Xi Zhang, Phillip A. Richmond, Bhavi P. Modi, Solenne Correard, Marius Gheorghe, Damir Baranašić, Walter Santana-Garcia, Ge Tan, Jeanne Cheneby, Benoit Ballester, Francois Parcy, Albin Sandelin, Boris Lenhard, Wyeth W. Wasserman, and Anthony Mathelier. JASPAR 2020: update of the open-access database of transcription factor binding profiles. Nucleic Acids Research, 48(D1):D87–D92, January 2020.

[16] Marius Gheorghe, Geir Kjetil Sandve, Aziz Khan, Jeanne Cheneby, Benoit Ballester, and Anthony Mathelier. A map of direct TF-DNA interactions in the human genome. Nucleic Acids Research, 47(4):e21, February 2019.

[17] Amirata Ghorbani, Abubakar Abid, and James Zou. Interpretation of Neural Networks Is Fragile. Proceedings of the AAAI Conference on Artificial Intelligence, 33(01):3681–3688, July 2019. Number: 01.

[18] Charles E. Grant, Timothy L. Bailey, and William Stafford Noble. FIMO: scanning for occurrences of a given motif. Bioinformatics, 27(7):1017–1018, April 2011.

[19] Sven Heinz, Christopher Benner, Nathanael Spann, Eric Bertolino, Yin C. Lin, Peter Laslo, Jason X. Cheng, Cornelis Murre, Harinder Singh, and Christopher K. Glass. Simple combinations of lineage-determining transcription factors prime cis-regulatory elements required for macrophage and B cell identities. Molecular Cell, 38(4):576–589, May 2010.

[20] Lukasz Huminiecki and Jaroslaw Horbanćzuk. Can We Predict Gene Expression by Understanding Proximal Promoter Architecture? Trends in Biotechnology, 0(0), April 2017.

[21] Arttu Jolma, Jian Yan, Thomas Whitington, Jarkko Toivonen, Kazuhiro R. Nitta, Pasi Rastas, Ekaterina Morgunova, Martin Enge, Mikko Taipale, Gonghong Wei, Kimmo Palin, Juan M. Vaquerizas, Renaud Vincentelli, Nicholas M. Luscombe, Timothy R. Hughes, Patrick Lemaire, Esko Ukkonen, Teemu Kivioja, and Jussi Taipale. DNA-Binding Specificities of Human Transcription Factors. Cell, 152(1–2):327–339, January 2013.

[22] Arttu Jolma, Yimeng Yin, Kazuhiro R. Nitta, Kashyap Dave, Alexander Popov, Minna Taipale, Martin Enge, Teemu Kivioja, Ekaterina Morgunova, and Jussi Taipale. DNA-dependent formation of transcription factor pairs alters their binding specificity. Nature, 527(7578):384–388, November 2015.

[23] Vineela Kadiyala, Sarah K Sasse, Mohammed O Altonsy, Reena Berman, Hong W Chu, Tzu L Phang, and Anthony N Gerber. Cistrome-based cooperation between airway epithelial glucocorticoid receptor and NF-κB orchestrates anti-inflammatory effects. Journal of Biological Chemistry, 291(24):12673–12687, 2016. Publisher: ASBMB.

[24] David R. Kelley, Yakir A. Reshef, Maxwell Bileschi, David Belanger, Cory Y. McLean, and Jasper Snoek. Sequential regulatory activity prediction across chromosomes with convolutional neural networks. Genome Research, 28(5):739–750, May 2018.

[25] Semyon Kolmykov, Ivan Yevshin, Mikhail Kulyashov, Ruslan Sharipov, Yury Kondrakhin, Vsevolod J Makeev, Ivan V Kulakovskiy, Alexander Kel, and Fedor Kolpakov. GTRD: an integrated view of transcription regulation. Nucleic Acids Research, 49(D1):D104–D111, January 2021.

[26] Peter K. Koo and Sean R. Eddy. Representation learning of genomic sequence motifs with convolutional neural networks. PLOS Computational Biology, 15(12):e1007560, December 2019.

[27] Judith F. Kribelbauer, Chaitanya Rastogi, Harmen J. Bussemaker, and Richard S. Mann. Low-Affinity Binding Sites and the Transcription Factor Specificity Paradox in Eukaryotes. Annual Review of Cell and Developmental Biology, 35(1):357–379, October 2019.

[28] Ivan V. Kulakovskiy, Ilya E. Vorontsov, Ivan S. Yevshin, Ruslan N. Sharipov, Alla D. Fedorova, Eugene I. Rumynskiy, Yulia A. Medvedeva, Arturo Magana-Mora, Vladimir B. Bajic, Dmitry A. Papatsenko, Fedor A. Kolpakov, and Vsevolod J. Makeev. HOCOMOCO: towards a complete collection of transcription factor binding models for human and mouse via large-scale ChIP-Seq analysis. Nucleic Acids Research, 2017.

[29] Marina Kulik, Melissa Bothe, Gözde Kibar, Alisa Fuchs, Stefanie Schöne, Stefan Prekovic, Isabel Mayayo-Peralta, Ho-Ryun Chung, Wilbert Zwart, Christine Helsen, Frank Claessens, and Sebastiaan H Meijsing. Androgen and glucocorticoid receptor direct distinct transcriptional programs by receptor-specific and shared DNA binding sites. Nucleic acids research, 49(7):3856–3875, 2021. Publisher: Oxford University Press.

[30] Michal Levo, Einat Zalckvar, Eilon Sharon, Ana Carolina Dantas Machado, Yael Kalma, Maya Lotam-Pompan, Adina Weinberger, Zohar Yakhini, Remo Rohs, and Eran Segal. Unraveling determinants of transcription factor binding outside the core binding site. Genome Research, 25(7):1018–1029, January 2015.

[31] Ming Li, Bin Ma, and Lusheng Wang. Finding similar regions in many strings. In Proceedings of the thirty-first annual ACM symposium on Theory of Computing, STOC ‘99, pages 473–482, New York, NY, USA, May 1999. Association for Computing Machinery.

[32] Christophe Menichelli, Vincent Guitard, Rafael M. Martins, Sophie Lebre, Jose-Juan Lopez-Rubio, Charles-Henri Lecellier, and Laurent Bréhélin. Identification of long regulatory elements in the genome of Plasmodium falciparum and other eukaryotes. PLOS Computational Biology, 17(4):e1008909, April 2021. Publisher: Public Library of Science.

[33] Leonid A. Mirny. Nucleosome-mediated cooperativity between transcription factors. Proceedings of the National Academy of Sciences of the United States of America, 107(52):22534–22539, December 2010.

[34] Ekaterina Morgunova and Jussi Taipale. Structural perspective of cooperative transcription factor binding. Current Opinion in Structural Biology, 47:1–8, December 2017.

[35] Gherman Novakovsky, Manu Saraswat, Oriol Fornes, Sara Mostafavi, and Wyeth W. Wasserman. Biologically relevant transfer learning improves transcription factor binding prediction. Genome Biology, 22(1):280, September 2021.

[36] Daniel Quang and Xiaohui Xie. DanQ: a hybrid convolutional and recurrent deep neural network for quantifying the function of DNA sequences. Nucleic Acids Research, 44(11):e107–e107, June 2016.

[37] Franziska Reiter, Sebastian Wienerroither, and Alexander Stark. Combinatorial function of transcription factors and cofactors. Current Opinion in Genetics & Development, 43:73–81, April 2017.

[38] Shuxiang Ruan and Gary D. Stormo. Inherent limitations of probabilistic models for protein-DNA binding specificity. PLOS Computational Biology, 13(7):e1005638, July 2017. Publisher: Public Library of Science.

[39] Shuxiang Ruan and Gary D. Stormo. Comparison of discriminative motif optimization using matrix and DNA shape-based models. BMC Bioinformatics, 19(1):86, March 2018.

[40] T D Schneider and R M Stephens. Sequence logos: a new way to display consensus sequences. Nucleic Acids Research, 18(20):6097–6100, October 1990.

[41] Tesa M Severson, Yongsoo Kim, Stacey EP Joosten, Karianne Schuurman, Petra Van Der Groep, Cathy B Moelans, Natalie D Ter Hoeve, Quirine F Manson, John W Martens, Carolien HM Van Deurzen, Ellis Barbe, Ingrid Hedenfalk, Peter Bult, Vincent T. H. B. M. Smit, Sabine C. Linn, Paul J. van Diest, Lodewyk Wessels, and Wilbert Zwart. Characterizing steroid hormone receptor chromatin binding landscapes in male and female breast cancer. Nature communications, 9(1):1–12, 2018. Publisher: Nature Publishing Group.

[42] Ning Shen, Jingkang Zhao, Joshua L. Schipper, Yuning Zhang, Tristan Bepler, Dan Leehr, John Bradley, John Horton, Hilmar Lapp, and Raluca Gordan. Divergence in DNA Specificity among Paralogous Transcription Factors Contributes to Their Differential In Vivo Binding. Cell Systems, 6(4):470–483.e8, April 2018.

[43] Richard I. Sherwood, Tatsunori Hashimoto, Charles W. O’Donnell, Sophia Lewis, Amira A. Barkal, John Peter van Hoff, Vivek Karun, Tommi Jaakkola, and David K. Gifford. Discovery of directional and nondirectional pioneer transcription factors by modeling DNase profile magnitude and shape. Nature Biotechnology, 32(2):171–178, February 2014.

[44] Divyanshi Srivastava and Shaun Mahony. Sequence and chromatin determinants of transcription factor binding and the establishment of cell type-specific binding patterns. Biochimica et Biophysica Acta (BBA) - Gene Regulatory Mechanisms, 1863(6):194443, June 2020.

[45] Robert E. Thurman, Eric Rynes, Richard Humbert, Jeff Vierstra, Matthew T. Maurano, Eric Haugen, Nathan C. Sheffield, Andrew B. Stergachis, Hao Wang, Benjamin Vernot, Kavita Garg, Sam John, Richard Sandstrom, Daniel Bates, Lisa Boatman, Theresa K. Canfield, Morgan Diegel, Douglas Dunn, Abigail K. Ebersol, Tristan Frum, Erika Giste, Audra K. Johnson, Ericka M. Johnson, Tanya Kutyavin, Bryan Lajoie, Bum-Kyu Lee, Kristen Lee, Darin London, Dimitra Lotakis, Shane Neph, Fidencio Neri, Eric D. Nguyen, Hongzhu Qu, Alex P. Reynolds, Vaughn Roach, Alexias Safi, Minerva E. Sanchez, Amartya Sanyal, Anthony Shafer, Jeremy M. Simon, Lingyun Song, Shinny Vong, Molly Weaver, Yongqi Yan, Zhancheng Zhang, Zhuzhu Zhang, Boris Lenhard, Muneesh Tewari, Michael O. Dorschner, R. Scott Hansen, Patrick A. Navas, George Stamatoyannopoulos, Vishwanath R. Iyer, Jason D. Lieb, Shamil R. Sunyaev, Joshua M. Akey, Peter J. Sabo, Rajinder Kaul, Terrence S. Furey, Job Dekker, Gregory E. Crawford, and John A. Stamatoyannopoulos. The accessible chromatin landscape of the human genome. Nature, 489(7414):75–82, September 2012.

[46] Robert Tibshirani. Regression Shrinkage and Selection Via the Lasso. Journal of the Royal Statistical Society, Series B, 58:267–288, 1994.

[47] Jimmy Vandel, Océane Cassan, Sophie Lebre, Charles-Henri Lecellier, and Laurent Bréhélin. Probing transcription factor combinatorics in different promoter classes and in enhancers. BMC Genomics, 20(1):103, February 2019.

[48] Jie Wang, Jiali Zhuang, Sowmya Iyer, XinYing Lin, Troy W. Whitfield, Melissa C. Greven, Brian G. Pierce, Xianjun Dong, Anshul Kundaje, Yong Cheng, Oliver J. Rando, Ewan Birney, Richard M. Myers, William S. Noble, Michael Snyder, and Zhiping Weng. Sequence features and chromatin structure around the genomic regions bound by 119 human transcription factors. Genome Research, 22(9):1798–1812, January 2012. Company: Cold Spring Harbor Laboratory Press Distributor: Cold Spring Harbor Laboratory Press Institution: Cold Spring Harbor Laboratory Press Label: Cold Spring Harbor Laboratory Press Publisher: Cold Spring Harbor Lab.

[49] Wyeth W. Wasserman and Albin Sandelin. Applied bioinformatics for the identification of regulatory elements. Nature Reviews Genetics, 5(4):276–287, April 2004.

[50] John W. Whitaker, Zhao Chen, and Wei Wang. Predicting the Human Epigenome from DNA Motifs. Nature methods, 12(3):265–272, March 2015.

[51] Rebecca Worsley Hunt, Anthony Mathelier, Luis del Peso, and Wyeth W. Wasserman. Improving analysis of transcription factor binding sites within ChIP-Seq data based on topological motif enrichment. BMC Genomics, 15(1):472, June 2014.

[52] Zeba Wunderlich and Leonid A. Mirny. Different gene regulation strategies revealed by analysis of binding motifs. Trends in genetics: TIG, 25(10):434–440, October 2009.

[53] An Zheng, Michael Lamkin, Hanqing Zhao, Cynthia Wu, Hao Su, and Melissa Gymrek. Deep neural networks identify sequence context features predictive of transcription factor binding. Nature machine intelligence, 3(2):172–180, February 2021.

[54] Jian Zhou and Olga G. Troyanskaya. Predicting effects of noncoding variants with deep learning-based sequence model. Nature Methods, 12(10):931–934, October 2015.

