## Supplementary Figures for "Systematic analysis of the genomic features involved in the binding preferences of transcription factors"

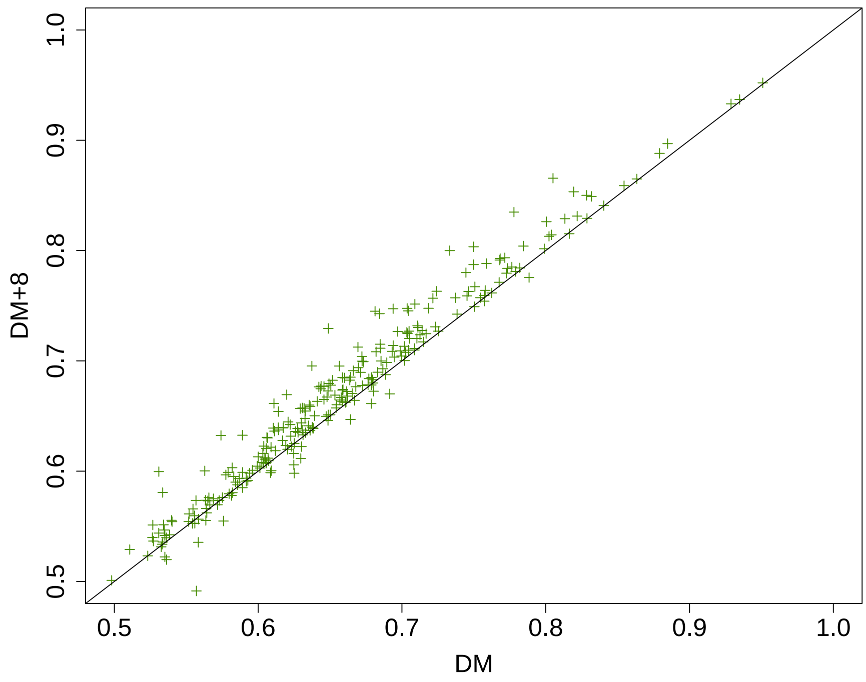


**Supp. Fig. 1**: **Increasing PWM length improves DM accuracy.** This dotplot compares the AUROC achieved by the DM model with length equal to that of the original JASPAR PWM and the DM model with 4 supplementary nucleotides on both sides of the PWM (DM+8).

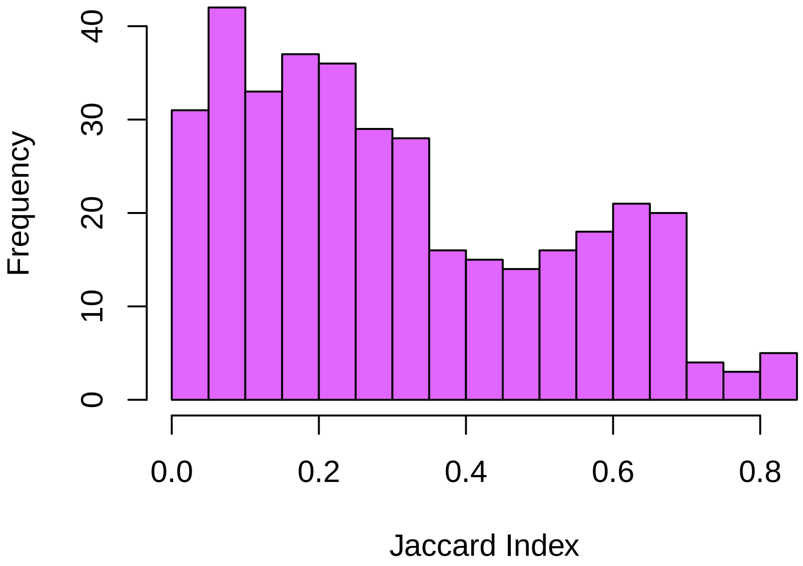


**Supp. Fig. 2**: **Distribution of Jaccard indexes of the 272 pairs of ChIP-seq experiments targeting a common TF in two different cell types.** The Jaccard index measures the proportion of peaks common to two ChIP-seq experiments, with regard to the total number of different peaks in the two experiments, *i.e.* Jaccard index = $\frac{|intersection|}{|union|}$.

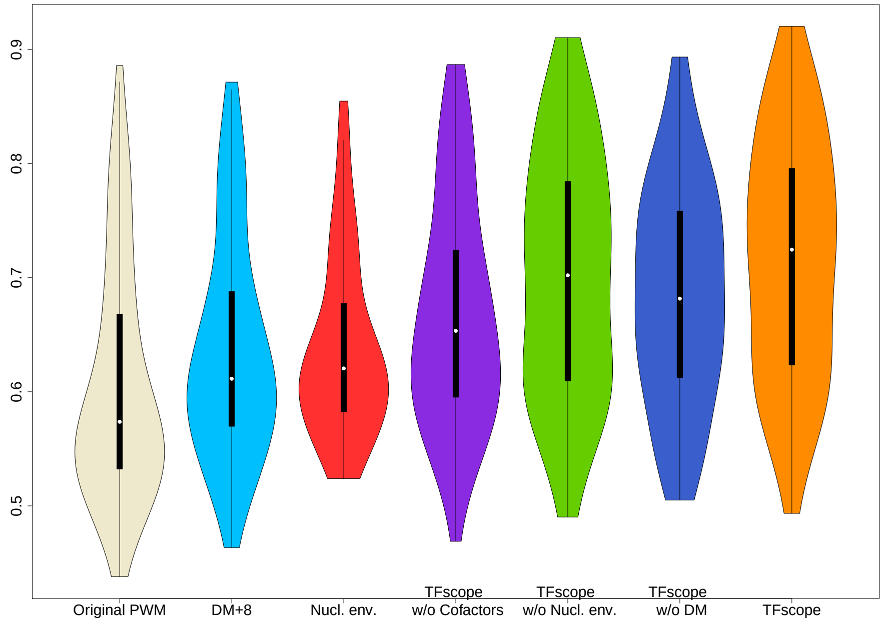


**A**

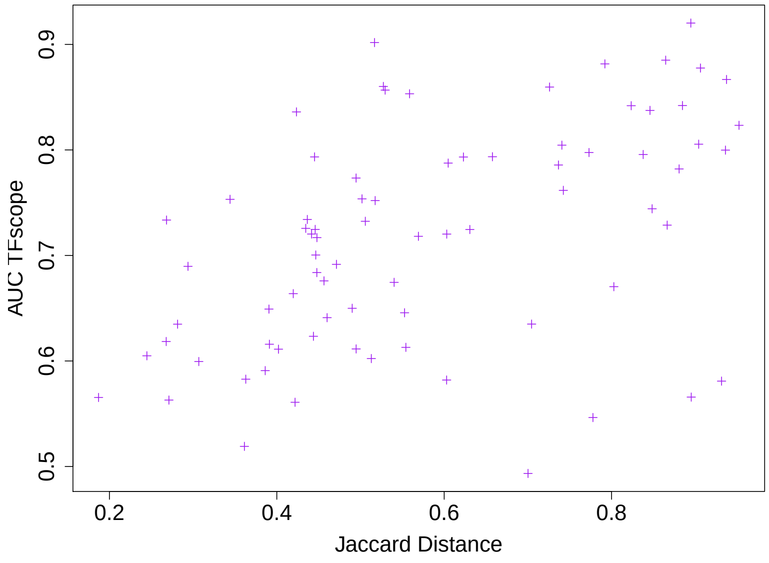


**B**

**Supp. Fig. 3**: **Discriminating binding sites of different treatments.** **A** Distribution of AUROCs achieved by TFscope and several alternative models for discriminating binding sites of one TF in two different treatments. **B** Link between TFscope accuracy and the similarity of ChIP-seq peaks in the two treatments. ChIP-seq experiments that have high proportion of peaks in common have low Jaccard distance. Pearson correlation = 0.48

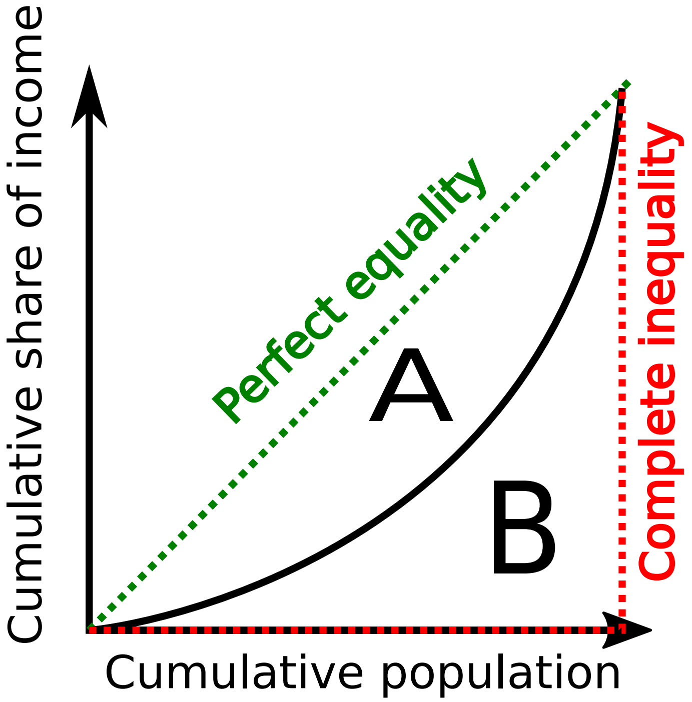


**Supp. Fig. 4**: **Lorenz curve and Gini coeficient**
